# Scout-based Multi-Echo NAvigator (SMENA) for high temporal resolution motion and B_0_ estimation and correction: applications to multi-echo GRE and EPTI

**DOI:** 10.64898/2026.06.10.731422

**Authors:** Nan Wang, Yimeng Lin, Yannick Brackenier, Aizada Nurdinova, Zihan Zhou, Daniel Abraham, Xiaozhi Cao, Congyu Liao, Kawin Setsompop

**Affiliations:** Department of Radiology, Stanford University, Stanford, CA, United States; Department of Electrical Engineering, Stanford University, Stanford, CA, United States; Department of Biomedical Engineering, School of Biomedical Engineering and Imaging Sciences, King’s College London, London, United Kingdom; GE Healthcare, MR Research China, Beijing, China; Neuroimaging Technology Research Center, Department of Radiology, University of California, San Francisco, CA, United States; Cognitive and Neurobiological Imaging Center, Stanford University, Stanford, CA, United States, 94305

**Keywords:** motion, B_0_ perturbation, data-driven estimation, high-temporal-resolution estimation

## Abstract

**Purpose:** To develop a data-driven technique, Scout-based Multi-Echo NAvigator (SMENA), for joint estimation of motion and B_0_ inhomogeneity (*δ*B_0_) at a temporal resolution of ∼200 ms with minimal additional scan time for gradient-echo acquisition.

**Methods:** SMENA consists of two key acquisition components: SMENA-scout and SMENA-nav. SMENA-scout is a rapid 3D 4-mm multi-echo acquisition completed in less than 8 seconds, providing images with matched contrast and phase at multiple echo times. SMENA-nav captures signal variations induced by motion and *δ*B_0_ during the scan using compact multi-echo navigator trajectories (3.5 ms) embedded within each TR. Motion and *δ*B_0_ maps were jointly estimated every ∼200 ms through a model-based optimization framework relating SMENA-scout to SMENA-nav. The estimation accuracy and correction performance of SMENA were evaluated in simulations and in vivo using multi-echo GRE and GRE-EPTI acquisitions. Multiple prospective motion experiments, including large continuous movement and deep breathing, were investigated.

**Results:** In both simulations and in vivo experiments, accurate motion and *δ*B_0_ estimation were achieved. Compared with motion-only estimation, joint estimation reduced rotation and translation errors. Joint motion and *δ*B_0_ correction resulted in substantial improvements in image quality, particularly at longer echo times, producing an NRMSE of 10.4% compared to 31.6% with motion-only correction. High-temporal-resolution tracking of motion and *δ*B_0_ enabled improved reconstruction quality in scenarios involving continuous motion and deep breathing.

**Conclusion:** SMENA enables high-temporal-resolution joint estimation of motion and *δ*B_0_ with minimal additional acquisition cost, providing a practical solution for motion- and *δ*B_0_-robust MRI.

## 1. Introduction

Patient bulk motion and fluctuation in the magnetic field inhomogeneity (B_0_) are major sources of image artifacts in MRI^1–6^. In brain imaging, bulk motion is typically approximated as rigid motion consisting of translations and rotations. Most MRI acquisitions are sensitive to such motion, leading to degraded image quality or prolonged scan times due to the need to repeat acquisitions. In addition, many sequences are also sensitive to *δ*B_0_ perturbations (*δ*B_0_). Sequences with long echo time (TE) and/or long readouts, such as multi-echo GRE and multi-shot Echo Planar Imaging (EPI), are particularly vulnerable to *δ*B_0_, resulting in ghosting and blurring artifacts^3,4^. Bulk motion and *δ*B_0_ are closely related, as motion is a major contributor to dynamic B_0_ variation. Physiological processes and system drift can further introduce additional B_0_ variations. Therefore, accurate knowledge of both motion and *δ*B_0_ during the scan is essential for effective artifact correction.

Over the years, a wide range of approaches have been proposed to estimate motion and/or *δ*B_0_. These methods are broadly categorized into four groups. The first category uses external devices^7–11^, such as optical devices and pilot tone approaches. These methods usually provide motion tracking at high temporal resolution (<100 ms), and some can estimate rigid motion parameters in real time, enabling prospective motion correction while preserving optimal k-space sampling^12^. However, these approaches require additional hardware and calibration, and tracking *δ*B_0_ remains challenging. This limitation becomes more pronounced at ultrahigh field strengths, where even small motions can induce substantial B_0_ changes.

The second category incorporates low-resolution volumetric navigator (vNAV) blocks into the acquisition. Examples include single-or dual-echo EPI^13,14^, stack-of-spiral^15^, 3D spiral^16^, FatNav^17–20^, and Echo Planar Time-Resolved Imaging (EPTI)^21^. These navigators enable robust image-domain estimation of motion and *δ*B_O_. However, vNAVs are typically inserted during sequence dead time or, if no dead time is available, incorporated as separate navigator modules at the cost of increased scan time. Nonetheless, either approach limits the temporal resolution of tracking to several seconds.

The third category employs self-navigating trajectories, such as PROPELLER^22^, MOJITO^23^, and DISORDER^24^. These trajectories are designed to repeatedly sample the center region of k-space to obtain low-spatial-frequency information for motion and *δ*B_0_ estimation.

While effective, they require specialized trajectory designs and are constrained by limited temporal resolution dictated by the sampling scheme.

The fourth category estimates motion and/or *δ*B_0_ utilizing compact navigators that traverse a limited region in k-space within a few milliseconds and can be inserted before or after each acquisition block. These approaches enable high-temporal-resolution estimation on order of TR. One subcategory directly estimates motion and *δ*B_0_ from signal variations in the navigator data^25,26^. These navigators are designed to move along lines and arcs in the k-space to capture motion-induced changes. However, their accuracy can degrade under large out-of-plane motion. Recently developed Servo Navigators^27–29^ have demonstrated accurate estimation of motion and first-order *δ*B_0_ under small-to-moderate motion. Estimation of large motion and higher-order *δ*B_0_ estimation is yet to be demonstrated.

Within this category, another subcategory, scout-based navigators, leverages a set of scout images to guide the motion and/or *δ*B_0_ estimation from compact navigators. The scout images are low-resolution images acquired at the beginning of the imaging session, with similar contrast to the navigator data. The estimation is usually performed through solving a motion and/or *δ*B_0_ informed signal equation relating scout images to navigator data. Examples, including free induction decay (FID) navigators for either motion^30^ or *δ*B_0_^31^ estimation and SAMER^32,33^ for motion estimation, have shown promising performance. However, simultaneous estimation of both motion and *δ*B_0_ remains challenging. Along this direction, our group previously developed QUEEN^34^, which combines non-Cartesian SPINS navigators (3.5 ms) with quantitative scout images to enable joint estimation of motion and *δ*B_0_ during contrast-changing scans. While promising, QUEEN has not been demonstrated on challenging cases with large motion and B0 perturbations, and its non-Cartesian navigators are susceptible to B_0_ and gradient imperfections, potentially limiting estimation accuracy.

To address the abovementioned challenges and enable motion- and *δ*B_0_-robust neuroimaging, in this work, we developed **S**cout-based **M**ulti-**E**cho **NA**vigator (**SMENA**). Compared to existing techniques, the key innovations of SMENA include: (1) rapid acquisition of 3D multi-echo scout images (SMENA-scout) capturing both magnitude and phase evolution for around 5 to 8 seconds using the EPTI framework^35–37^; (2) a 3.5-ms multi-echo navigator (SMENA-nav) embedded in every TR; (3) joint estimation of motion and *δ*B_0_ at a temporal resolution of 200ms; and (4) flexible integration with gradient-echo sequences. For the proof-of-concept, SMENA was incorporated into conventional multi-echo GRE and more advanced GRE-EPTI^35–37^. Its ability to provide high-temporal-resolution tracking of motion and *δ*B_0_ was demonstrated across multiple in vivo experiments, showing robust and consistent performance across subjects.

## 2. Methods

### 2.1. Motion/ *δ*B_0_ estimation pipeline for scout-based navigators^32^

Conventional scout-based navigators estimate rigid motion or *δ*B_0_ parameters by solving the following nonlinear least-square problem

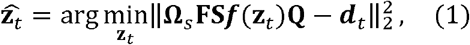

Where **Q** is the single-contrast scout image,**F** is the Fourier transform,**S** is the multi-channel coil sensitivity **Ω**_*s*_ is the compact navigator sampling pattern,*d* _*t*_ is the multi-channel navigator data at state *t* (*t*=1, …,*N*_*t*_ with *N*_*t*_ as the total number of states to be estimated),**z**_*t*_ is the motion or *δ*B_0_ parameters to be estimated at state *t* and *f*(·)is the corresponding motion or *δ*B_0_ operator. Because **Q** contains only single-contrast information and **Ω**_*s*_ samples a limited portion of k-space, simultaneous estimation of motion and *δ*B_0_ is inherently ill-conditioned.

### 2.2. SMENA-scout and SMENA-nav

To accurately decode the motion and *δ*B_0_ parameters from limited navigator data, the scout and navigator design must satisfy the following requirements: (1) both scout and navigator should capture magnitude and phase evolution; (2) for GRE sequences, the T1 and T2* contrast weightings as well as the baseline phase from B_0_ should be consistent between the scout image and the navigator data; (3) the scout acquisition should be completed within 10 seconds to minimize motion and *δ*B_0_ contaminations, while navigator duration should be below 4 ms to avoid disrupting the actual acquisition.

To fulfill these requirements, in this work, SMENA-scout utilizes a rapid GRE-EPTI^35–37^ acquisition (Figure 1A). It employs multi-echo Cartesian readout with k_y_-k_z_ blips to enable efficient spatiotemporal k-t space sampling. A low-rank subspace reconstruction was implemented to recover high-quality images with a high undersampling rate^37^. To ensure consistent T1 contrast between SMENA-scout and SMENA-nav, the TR and flip angle of SMENA-scout are matched to the actual acquisition. The T2* and B_0_ maps obtained from SMENA-scout allow the generation of reference images with consistent T2* weightings and baseline phase from B_0_ as of SMENA-nav. The spatial resolution for SMENA-scout is 4 mm, which is sufficient to correct motion and *δ*B_0_ for ∼0.5-mm acquisition^30–32^. The scan time of SMENA-scout scales proportionately with its TR. For an example TR of 50 ms, the scan time is 5.8 seconds.

**Figure 1:**
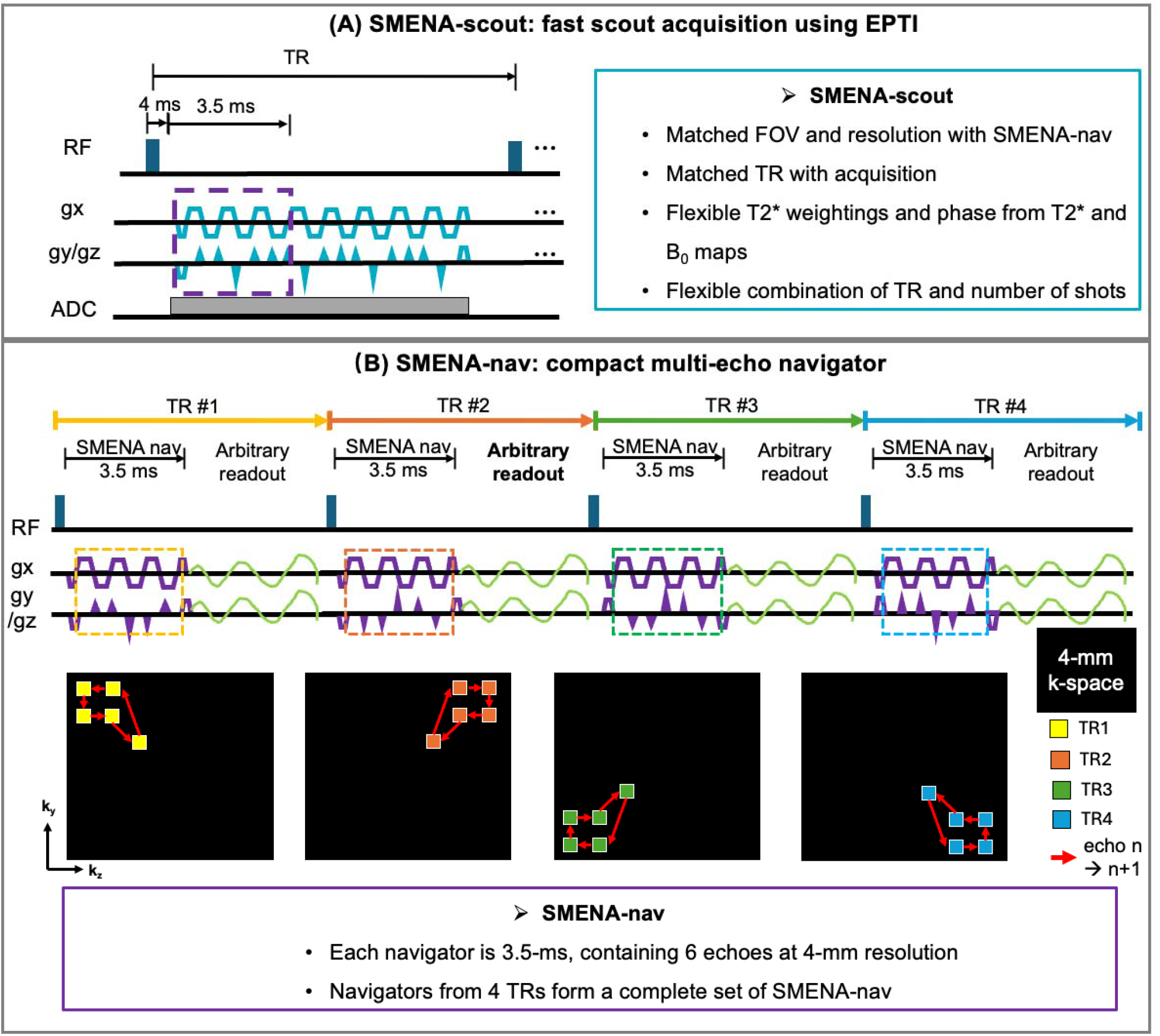
Demonstration of SMENA. (A) The strategy of SMENA-scout. EPTI trajectory is used for rapid acquisition of multi-echo scout images. The TR of SMENA-scout is set to be the same as the TR of the actual acquisition for a consistent T1 contrast between nav and scout; the images can be generated at any TE for consistent T2* and phase contrasts. (B) The design of SMENA-nav. SMENA-nav is a short 3.5-ms 6-echo Cartesian trajectory. The ky-kz locations within one navigator start from center, move to outter, and then back to center k-space, following the read arrows. The combination of navigators from 4 TRs provides one complete set for an estimation.

SMENA-nav is a 6-echo Cartesian trajectory with a duration of 3.5 ms, covering 4-mm extent in k_x_ (Figure 1B). Between echoes, there are k_y_-k_z_ blips to traverse to different k-space locations. Each navigator readout samples one quadrant of the 4-mm^2^ k_y_-k_z_ space to limit the duration and area of the blips for short echo spacing and low eddy-current effect. A complete navigator set is formed by combining data from four consecutive TRs, resulting in full coverage of the 4-mm k_y_-k_z_ space. Therefore, the temporal resolution of joint motion and *δ*B_0_ estimation is equal to four TRs (∼200 ms for a typical multi-echo GRE protocol at 3T with TR = 50 ms). Due to its short duration, SMENA-nav can be flexibly inserted within the sequence; in this work, it is placed at the beginning of each TR before the actual readouts.

To maximize sampling efficiency, aggressive ramp sampling is used, which introduces potential eddy current errors (Supporting Information Figure S1). To mitigate these errors, a field-correcting GRAPPA (FCG) technique^38,39^ was applied to both SMENA-scout and SMENA-nav, with details in Supporting Information Section A.

### 2.3. SMENA motion and *δ*B_0_ estimation pipeline

The motion and *δ*B_0_ is estimated through a model-based optimization framework relating SMENA-scout to SMENA-nav, as shown in Figure 2. The rigid motion is parameterized using six motion parameters (three for translation and three for rotation). For *δB*_0_ direct is estimated of a full 3D *δ*B_0_ map from limited navigator data is challenging. Given that *δ*B_0_ maps are spatially smooth^31^, they are modeled using a low-order spherical harmonic (SH) representation as*δB*_0_=**H**_***c***_, where **H** is the SH basis functions and is ***c*** the coefficients to be estimated. In this work, a second-order SH model is adopted following previous investigations at 3T^31,34^, resulting in 9 unknown coefficients. Taking together, the inverse problem for motion and *δ*B_0_ estimation in SMENA is formulated as

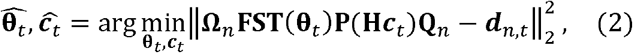

where **T(θ**_*t*_ )is the rigid motion operator^40^ with motion parameters **θ**_*t*_ at state *t*, 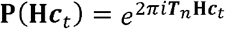 is the phase accumulation induced by *δB*_0_ across echo times ***T***_*n*_, **Ω**_*n*_ is the multi-echo sampling pattern of SMENA-nav across 4 TRs,**Q**_*n*_ is the multi-echo SMENA-scout, and *d*_*n,t*_ is the acquired multi-echo SMENA-nav data at state *t*. The unknown parameters are estimated iteratively using a quasi-Newton approach:

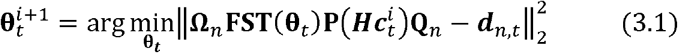

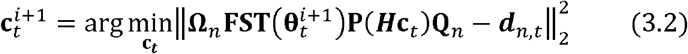

**Figure 2:**
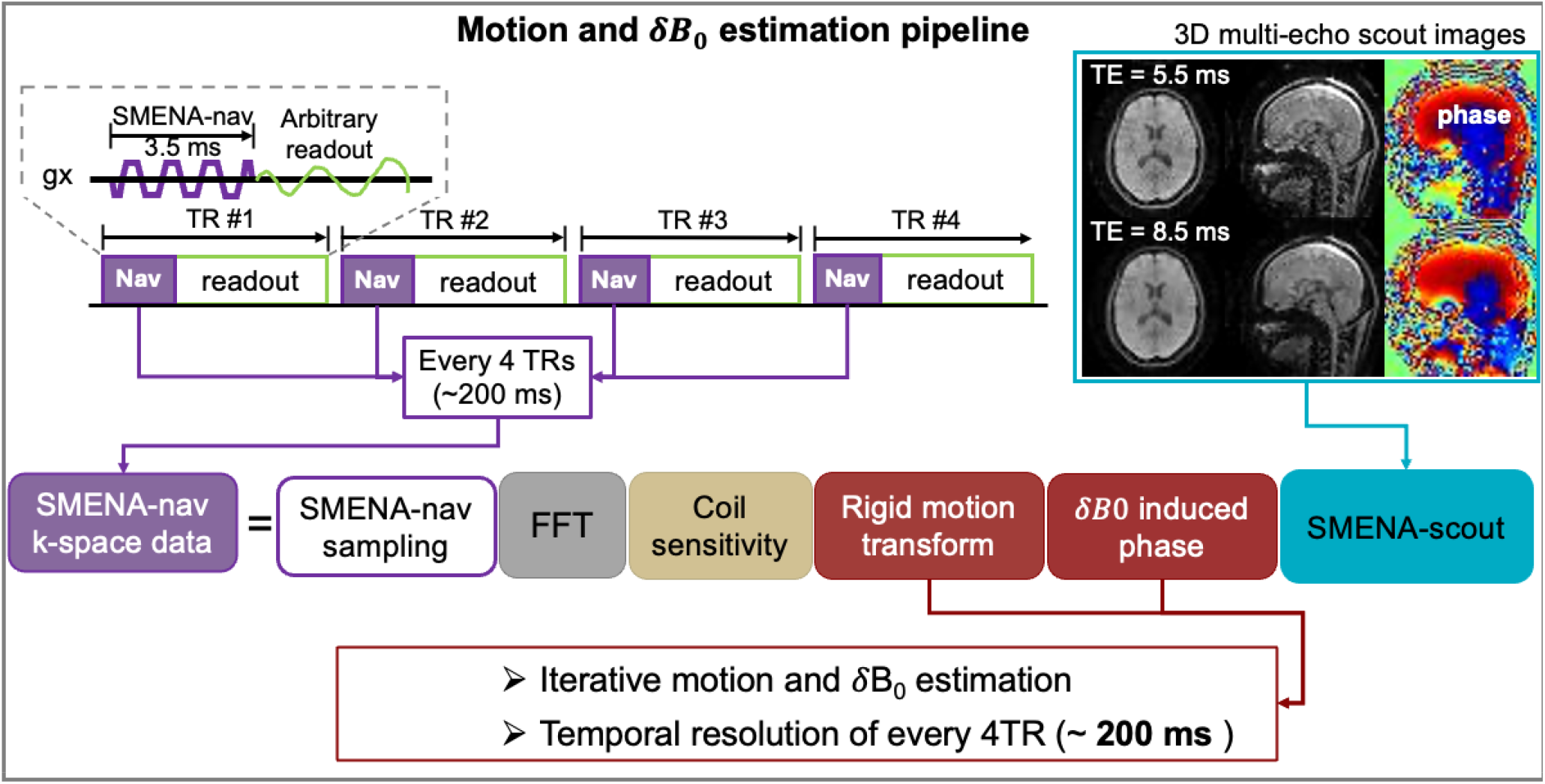
The pipeline for SMENA estimation. The navigators from every 4 TRs provide a set of motion and dB0 parameters by iteratively solving the optimization problem. The temporal resolution of motion and dB0 estimation is equal to the duration of 4 TRs. For a typical 3D multi-echo GRE, this is around 200 ms.

### 2.4. Optimization of the SMENA-nav sampling

The sensitivity of the navigator trajectory to motion and *δ*B_0_ depends on its k_y_-k_z_ locations. To evaluate this effect, the data consistency (DC) distance metric^33^ was used. Let *s*_*o,ij*_=**Ω**_*ij*_ **FSQ**_*n*_ denote the multi-channel data from the multi-echo SMENA-scout **Q**_*n*_ at location(*k*_*y, i*_,*k*_*z, j*_) at a reference state (**θ**_0_=**0**and**c**_o_=0),*s*_*r,ij*_= **Ω**_*ij*_ **FST** (θ_*r*_) **P**(***Hc***_*r*_ ***Q***_*n*_ denote the corresponding data after moving **Q**_*n*_ to a random motion/ *δ*B_0_ state with and **θ**_*r*_ and **c**_*r*,_ and *d* _*r,ij*_ denote the acquired multi-channel SMENA-nav data at the same location given the same motion/ *δ*B_0_ state withθ_*r*_ and **c**_*r*_. The DC difference between*s*_*0,ij*_ and *d*_*r,ij*_ is defined as

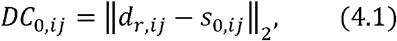

the DC difference between *s*_*r,ij*_, and *d*_*r,ij*_ is defined as

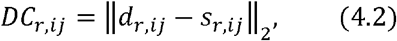

and the normalized DC distance metric at l ocati on (*k*_*y,i*_,*k*_*z*_,_*i*_) is then defined as

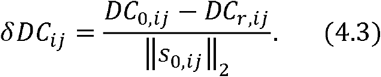

For a location(*k*_*y,i*_,*k*_*z,i*_) that provides high sensitivity to motion and *δ*B_0_ estimation,*DC*_*r, ij*_ it should be relatively small, and *DC*_*0,ij*_, should be sufficiently large, leading to a large *δDC*_*ij*_.In this work, *δDC*_*ij*_ was evaluated through simulations using multi-echo GRE data acquired at 3T from 20 subjects to ensure variability in anatomy and coil sensitivity. For each subject, 20 random motion/*δ*B_0_ states were generated. The final *δDC* map was obtained by averaging across all 400 simulations. Locations with larger *δDC* indicate higher sensitivity to motion and/or *δ*B_0_.

### 2.5. Motion- and *δ*B_0_-corrected reconstruction for multi-echo GRE and GRE-EPTI

For multi-echo GRE, the image reconstruction is performed for each echo separately. By incorporating state-specific encoding operators to account for motion and *δ*B_0_ variations, the per-echo reconstruction is formulated as

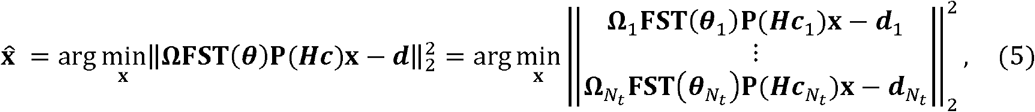

Where **x** is the image of a given echo time, **Ω**_*t*_ is the sampling pattern for multi-echo GRE at motion/*δ*B_0_ state *t*=1,…,*N*_*t*_ with *N*_*t*_ as the total number of motion and *δ*B_0_ states.

For EPTI, a low-rank subspace reconstruction^36,37^ incorporating motion and *δ*B_0_ is formulated in a similar manner:

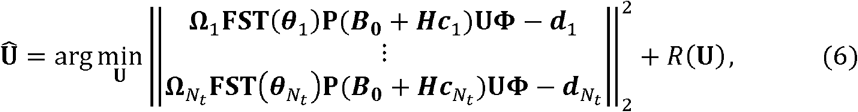

where ***B***_0_ is the baseline B_0_ map at the reference position (which is the same position as for the scout image). The baseline B_0_ map is estimated from a data-driven rank-shrinking B_0_ update algorithm^37^. To ensure a good quality, the motion and *δ*B_0_ correction are included in the rank-shrinking B_0_ update step as well.

The order of motion and B_0_ operators matters. In Equations 5 and 6, both the image to be solved and the phase induced by B_0_ and/or *δ*B_0_ are defined at the reference position, and the motion operator is applied afterward to reach the target position. An alternative formulation is to move the phase induced by B_0_/ *δ*B_0_ to the target position first and perform it on the motion-modulated images, as shown in the Supporting Information Equation S2. Nonetheless, this approach leads to elevated errors in image reconstruction, with details demonstrated in the Supporting Information Section B.

### 2.6. Simulations

Simulations were conducted to evaluate: (1) the estimation accuracy of joint motion and *δ*B_0_ estimation vs motion-only estimation; (2) the image quality improvement using joint motion and *δ*B_0_ correction vs motion-only correction.

The Simulation pipeline is demonstrated in Figure 3. Whole-brain multi-echo GRE images at 1-mm isotropic resolution were simulated from ground-truth (GT) parameter maps (including PD, T2*, B_0_, and sensitivity maps) acquired using a whole-brain 0.75-mm sEPTI^37^. The GT maps are higher in resolution than the target images to avoid k-space edge errors when performing rotation. Twelve sets of motion parameters and corresponding and *δ*B_0_ maps were generated using realistic parameters reported in previous studies^31^. The timing for the 12 motion/*δ*B_0_ states was randomly assigned to mimic realistic conditions. Taking together, a set of motion-and-*δ*B_0_ -corrupted 3D multi-echo GRE data at 1-mm resolution was simulated. The SMENA-nav were also simulated every TR. Besides, a 4-mm SMENA-scout acquisition was simulated from the same GT maps at the reference position.

**Figure 3:**
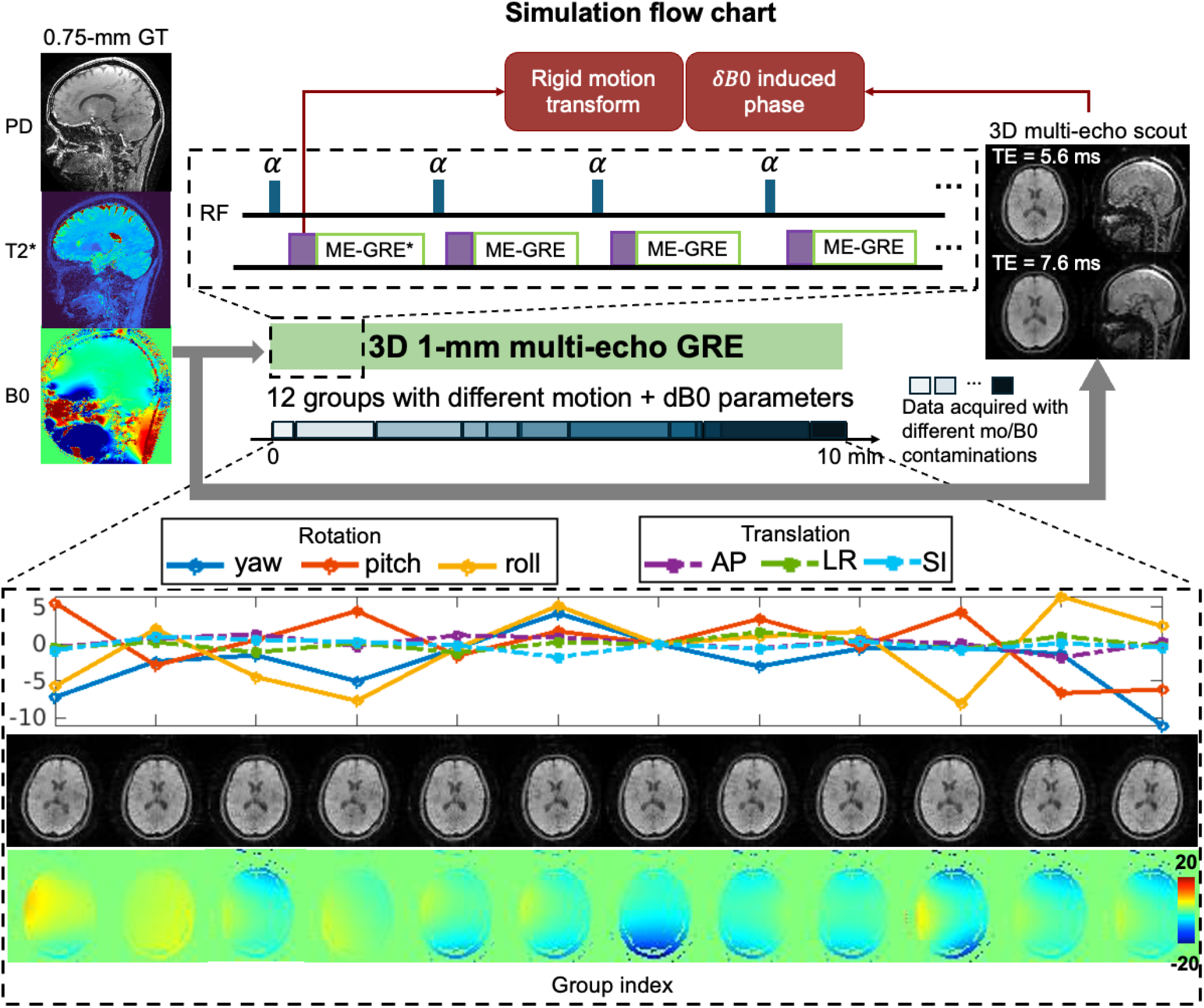
Simulations. Ground truth (GT) 3D 0.75-mm PD, T2*, B0, and sensitivity maps (not shown here) were modulated with 12 different motion and dB0 parameters. 3D 1-mm multi-echo GRE was simulated to mimic a motion-corrupted case, with blocks of different colors representing the data sampled from different motion/dB0 setups. The SMENA-nav was generated every TR along the simulations. The SMENA-scout was generated from GT images without motion and dB0 modulations, which is considered the reference position.

The accuracy of joint motion-and-*δ*B_0_ estimation and motion-only estimation was evaluated. In the reconstruction, the imaging data with no correction, motion-only correction, and joint motion-and-*δ*B_0_ correction were performed. The image quality was evaluated using the normalized root mean square error (NRMSE) with respect to GT images.

### 2.7. In vivo experiments

All in vivo experiments were approved by the institutional review board, with written informed consent obtained. Five subjects were included in the study on a 3T system (UHP, GE Healthcare, Milwaukee, USA). In each imaging session, the SMENA-scout (8 seconds) was acquired, followed by multiple scans using 3D whole-brain 1-mm EPTI with SMENA-nav embedded in each TR, as shown in Figure 4A and 4B. The first EPTI scan was acquired under normal breathing without voluntary motion, serving as a stationary reference. In the second EPTI scan, subjects performed deep breathing without voluntary motion, while mild involuntary motion (e.g., pitch rotation) could occur. In the third EPTI scan, subjects were asked to move 6-12 times at random time points, mimicking realistic motion patterns. Following all EPTI scans, a 3D whole-brain 1-mm isotropic multi-echo GRE was acquired, during which subjects were asked to perform 10-15 voluntary movements at random times.

**Figure 4:**
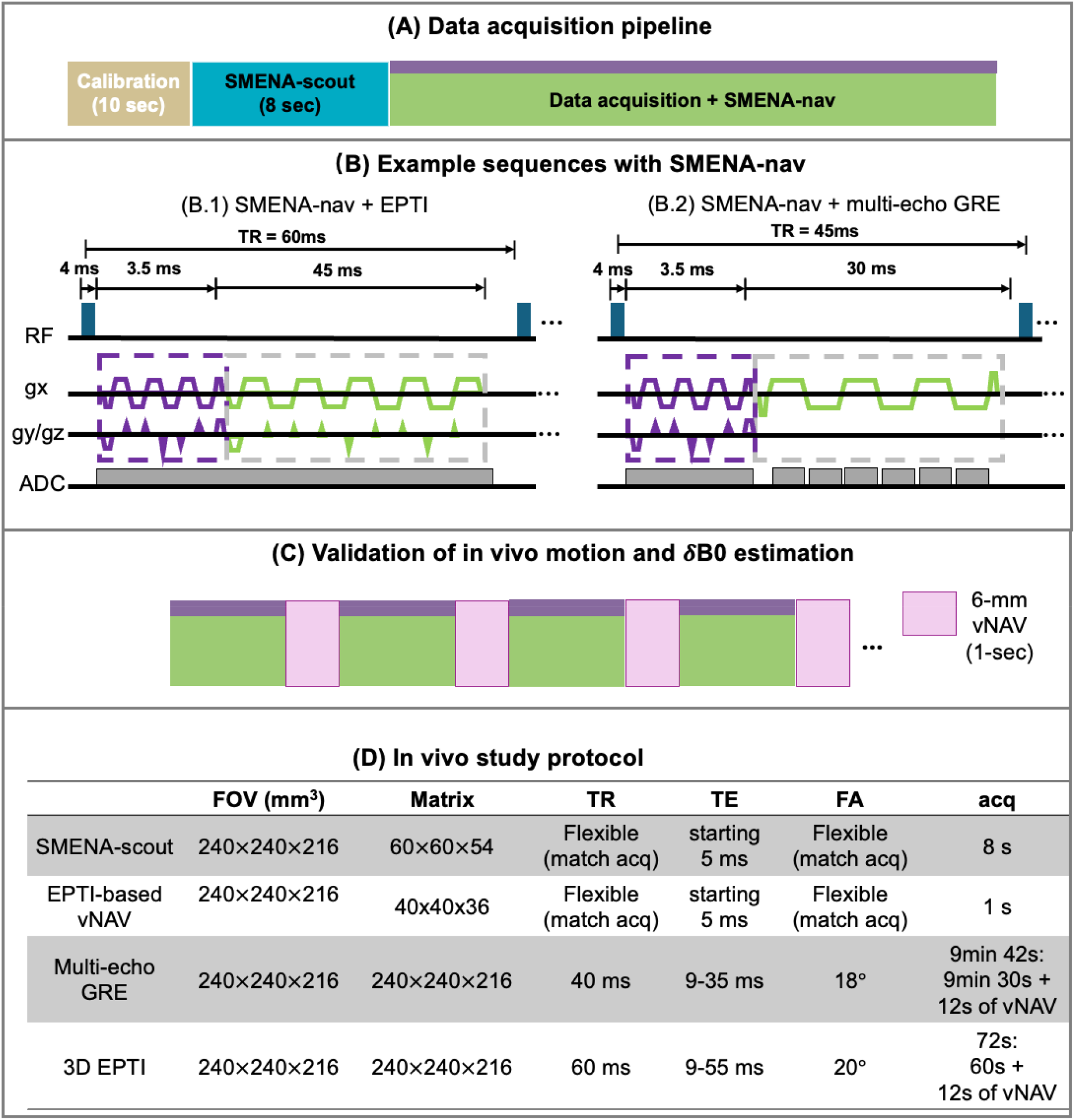
In vivo experiments. (A) Image acquisition pipeline with SMENA. Typically, a set of calibration data is acquired at the beginning to provide sensitivity maps, low-resolution B0, etc. Following this, SMENA-scout is acquired in 8 seconds. Then, sequences incorporated with SMENA-nav are performed as needed. (B) Example gradient waveforms for the incorporation of SMENA-nav and different types of readouts. (C) Demonstration of the insertion of vNAV during the data acquisition. The vNAV is a EPTI-based 6-mm low-resolution acquisition, which takes 1 second per volume. It serves the in vivo GT of motion and dB0 parameters at a few time points. (D) Detailed sequence parameters.

To obtain gold-standard motion and *δ*B_0_ measurements, a 6-mm EPTI-based vNAV^21^ was evenly inserted 12 times into each sequence, as demonstrated in Figure 4C. The acquisition time per vNAV is 1 second, leading to a 12-second increase in the scan time for each sequence. Image-based motion registration and *δ*B_0_ estimation was used. The temporal resolution of vNAV-based motion and *δ*B_0_ estimation is 5 seconds for EPTI and 50 seconds for multi-echo GRE. The detailed sequence parameters are listed in Figure 4D.

The motion and *δ*B_0_ parameters from SMENA estimation with TRs right after vNAV’s acquisition were used to compare with vNAV’s estimation. Image reconstruction was performed under four conditions: (1) no correction, (2) low-temporal-resolution joint motion-and-*δ*B_0_ correction using vNAV, (3) high-temporal-resolution motion-only correction using SMENA, and (4) high-temporal-resolution joint correction using SMENA.

## 3. Results

### 3.1. Optimization of the SMENA-nav sampling

The *δDC* map across the k_y_-k_z_ plane, averaged over the 400 simulated experiments, was evaluated for each echo of SMENA-nav. For the simplicity of display, Figure 5 shows representative *δDC* maps at echo 1 (TE = 5.6 ms), echo 3 (TE = 6.6 ms), and echo 5 (TE = 7.6 ms).

**Figure 5:**
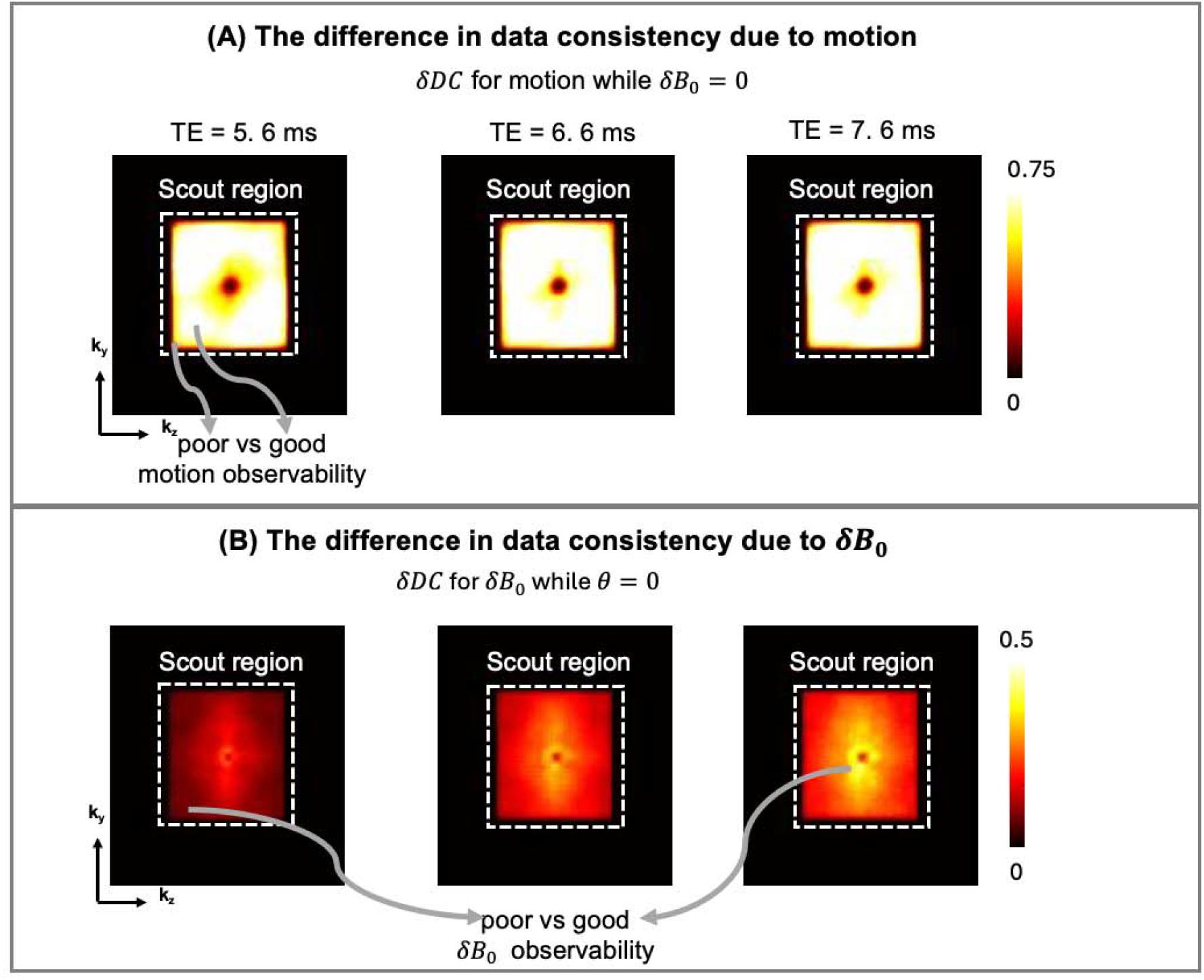
The results of the evaluation of DC distance. (A) For motion parameters, when there is motion, the *δDC* at a certain ky-kz location increase with the distance from center k-space. It decreases quickly when apporaching the edge of scout k-space (4 mm) due to edge errors. At different TE, *δDC* is similar. (B) For *δ*B0 parameters, *δDC* is large towards center k-space. It increases substantially with the increase of TE.

In the presence of motion, *δDC* is minimal near the center of k-space, increases toward intermediate spatial frequencies, and decreases near the edge of the 4-mm scout k-space support. This behavior is consistent with the physics of motion: translations pose linear phases on k-space, while rotations correspond to rotations of k-space coordinates. Both effects scale with distance from the k-space center, resulting in increased motion sensitivity at higher spatial frequencies; the decrease of the sensitivity towards the k-space edge is because the edge of the scout k-space presents errors due to its finite spatial resolution. The motion-induced *δDC* patterns are consistent across echo times.

For *δ*B_0_, the central k-space region (≤6 Δ*k*) excluding the very center (≤2 Δ*k*), exhibits higher*δDC* compared to outer k-space regions. This is consistent with the effect of*δ*B_0_ –induced phase, which manifests as a convolution in k-space. Due to higher SNR in the central k-space, *δ*B_0_ effects are more detectable in this region. Furthermore, *δ*B_0_ visibility increases with echo time, indicating that later echoes provide enhanced sensitivity to field variations.

Based on these observations, the SMENA-nav sampling pattern is designed such that k_y_-k_z_ locations across echoes start from central k-space, extend toward outer regions, and return to central k-space, thereby balancing sensitivity to both motion and *δ*B_0_.

### 3.2. Simulation results

The performance of SMENA for joint motion and *δ*B_0_ estimation is shown in Figure 6A. When performing motion-only estimation, the relative errors compared to GT are 3.8% for rotation and 17.1% for translation, averaged over 12 motion states. These errors are substantially reduced when performing joint estimation, yielding relative errors of 1.0% for rotation, 1.3% for translation, and 2.14% for *δ*B_0_ map averaged across all voxels.

**Figure 6:**
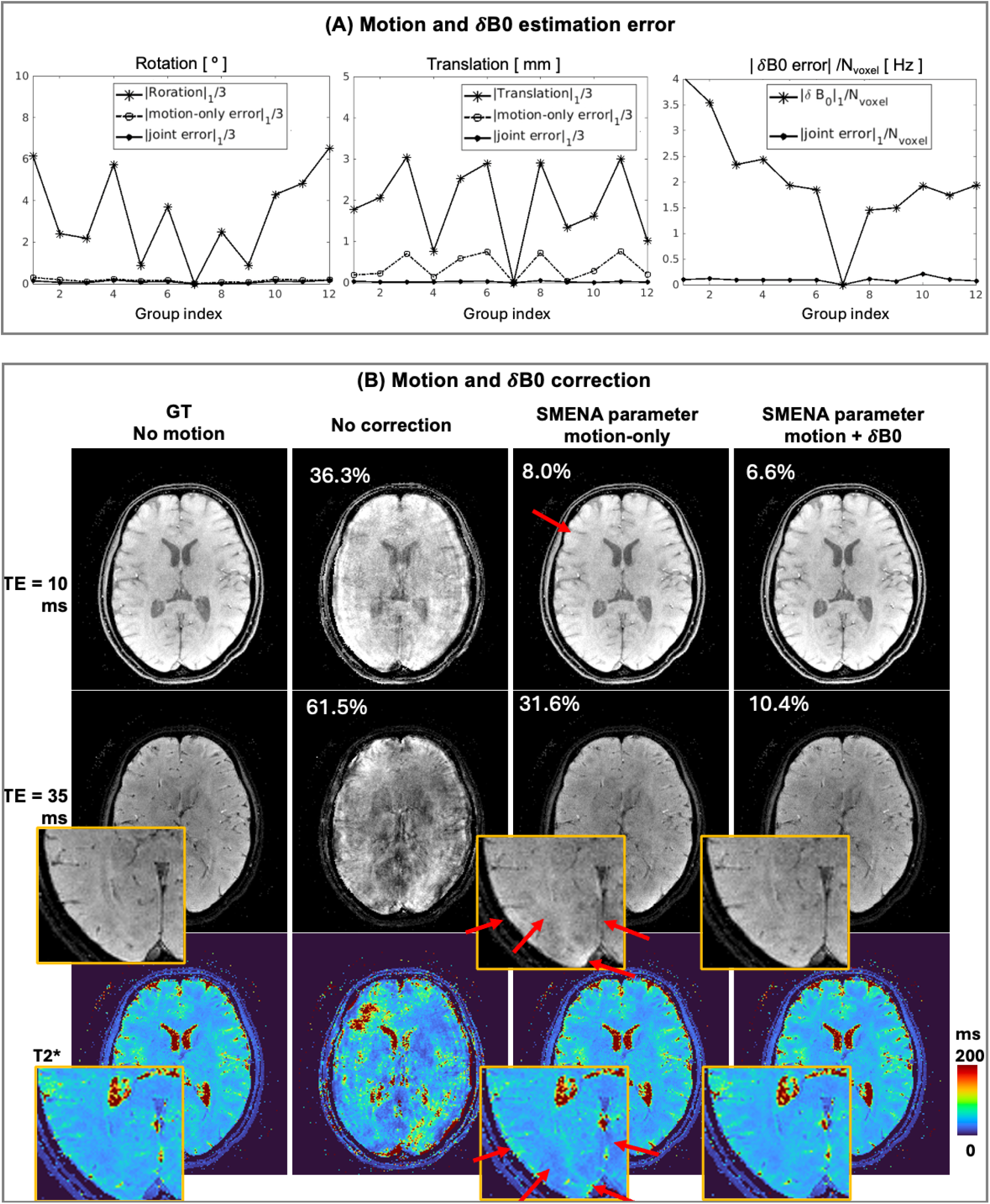
The results of the simulations. (A) for motion and *δ*B0 estimation, the motion-only estimation (dashed line with a hollow circle) possesses higher error than the joint motion+*δ*B0 estimation (solid line with solid circle), particularly for translation. The joint motion+*δ*B0 estimation shows low errors compared to GT. (B) the motion and *δ*B0 parameters estimated from SMENA can significantly improve the image quality in the reconstruction. For later TEs, joint motion+*δ*B0 correction significantly outperform motion-only correction. Some artifacts are labeled by red arrows.

In image reconstruction, substantial improvements are observed with joint correction. Without correction, motion-corrupted images exhibit NRMSE of 36.3% at TE = 10 ms and 61.5% at TE = 35 ms. Motion-only correction reduces NRMSE to 8.0% and 31.6% at the respective echo times. In contrast, joint motion-and-*δ*B_0_ correction further reduces NRMSE to 6.6% at TE = 10 ms and 10.4% at TE = 35 ms. Notably, the improvement is particularly pronounced at longer echo times, where B_0_ effects are stronger. Consistent trends are observed in T2* maps (Figure 6B), where joint correction yields the most accurate results.

### 3.3 In vivo results

Figure 7 shows in vivo joint motion-and-*δ*B_0_ estimation using SMENA in a multi-echo GRE experiment. In Figure 7A, motion estimates from SMENA (blue lines) demonstrate strong agreement with vNAV measurements (red circles) at the 12 vNAV sampling points. In addition, SMENA provides continuous motion tracking at a temporal resolution of 200 ms, revealing fine temporal dynamics that are not captured by vNAV. The corresponding *δ*B_0_ estimation is shown in Figure 7B. Example temporal evolution of 0th-, 1st-, and 2nd-order SH coefficients indicates that: (1) *δ*B_0_ are strongly correlated with bulk motion; and (2) oscillatory components at approximately 0.3 Hz are present, consistent with respiratory motion. At the vNAV sampling points, *δ*B_0_ maps estimated by SMENA are in good agreement with vNAV results. In contrast, motion-only estimation leads to noticeable errors, which is shown in Supporting Information Figure S3.

**Figure 7:**
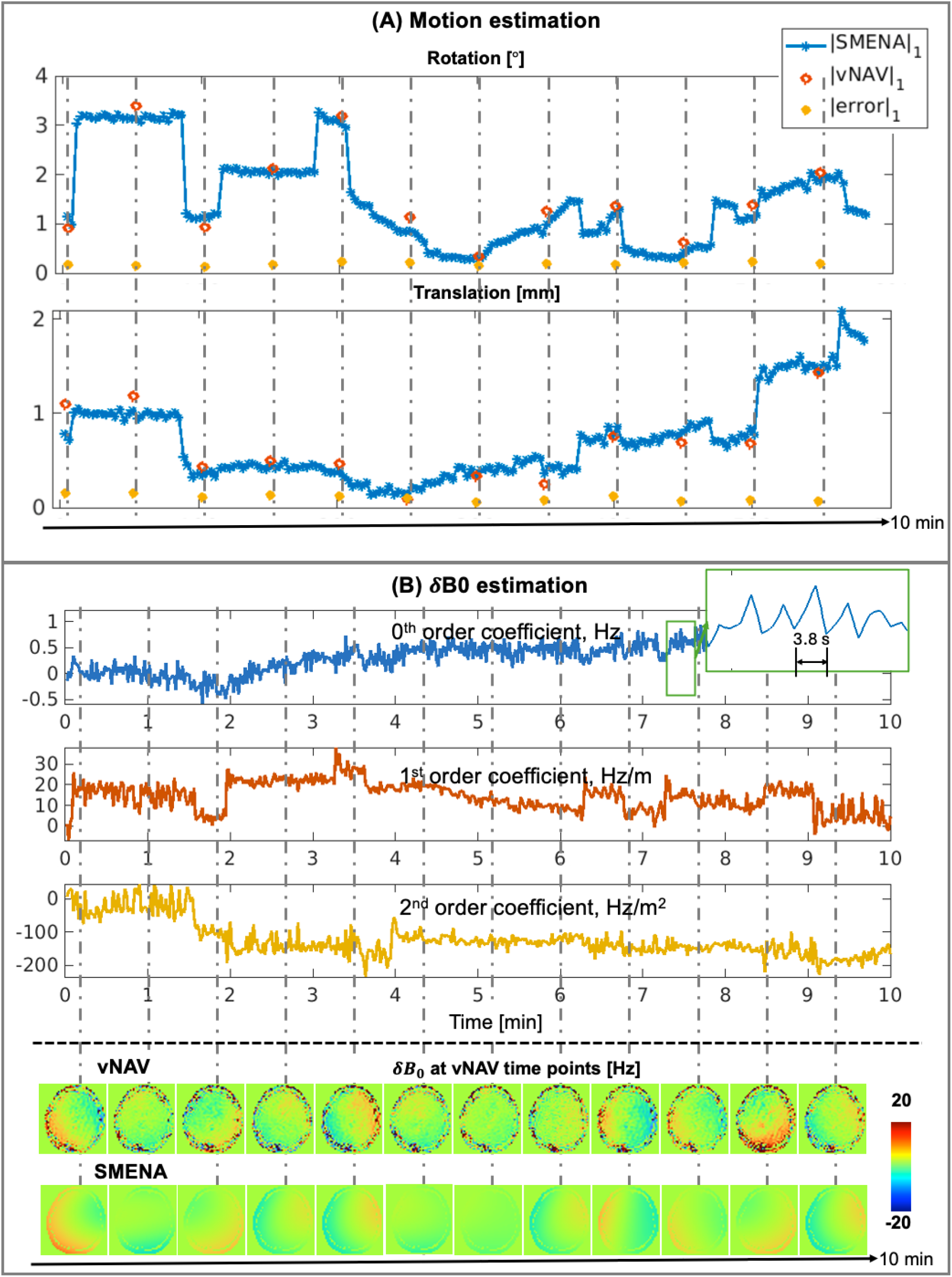
An in vivo example of SMENA motion and *δ*B0 estimation. Joint motion and *δ*B0 estimation were performed for all cases. (A) At the 12 discrete time points where vNAV is acquired, the motion parameters from SMENA’s joint estimation shows low errors compared to the motion parameters from vNAV. In the meantime, SMENA provides a nice temporal profile of the motion during the entire scan. (B) At the 12 discrete time points, the *δ*B0 maps from SMENA and vNAV are similar. In the meantime, the SMENA estimation provides much more temporal details. The oscillation along the example SH coefficients coincides with the frequency of respiration.

Figure 8 demonstrates the reconstruction performance for motion- and *δ*B_0_-corrupted multi-echo GRE data. Without correction, severe ringing artifacts are observed in multi-echo images, T2* maps, and QSM. Correction using vNAV parameters does not improve image quality due to its limited temporal resolution and inability to capture the correct timing of motion (as indicated by gray dashed lines in Figure 8A). Motion-only correction using SMENA’s estimation reduces artifacts in early echo images (TE = 10 ms), while significant artifacts remain in later echo images (TE = 40 ms) and in T2* and QSM maps due to uncorrected *δ*B_0_ effects. In contrast, joint high-temporal-resolution motion-and-*δ*B_0_ correction using SMENA produces the cleanest images and quantitative maps.

**Figure 8:**
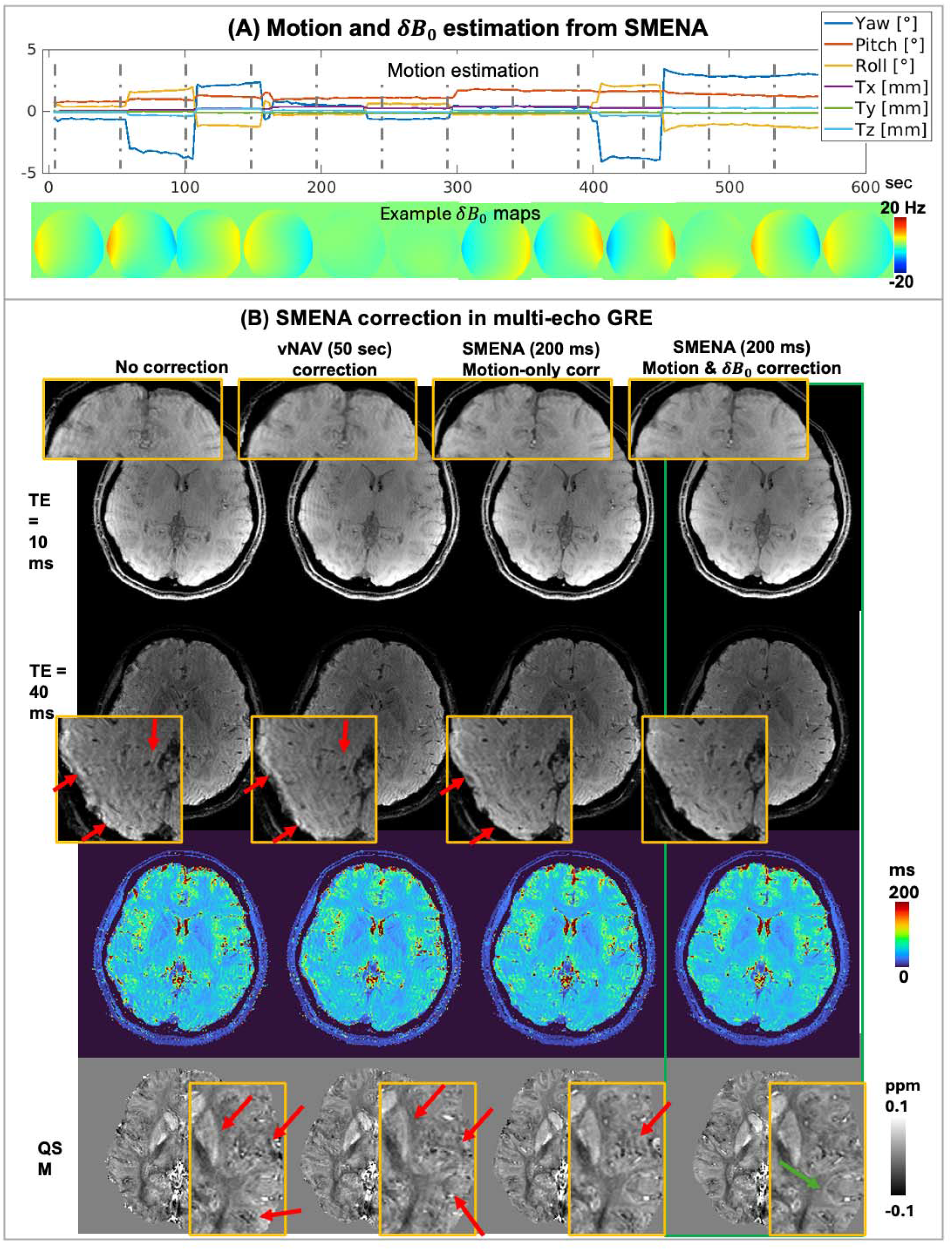
In vivo example of SMENA’s application to motion-corrupted multi-echo GRE. The low-temporal-resolution correction using the motion and *δ*B0 from vNAV every 50 seconds cannot provide good correction due to the wrong timing; the correction using SMENA’s estimation can significantly improve the image quality. The joint correction of motion and *δ*B0 provides the best results, which are labeled by green frames. The red arrows mark the example artifacts, while the green arrows mark the improved structures.

Figure 9 shows results for EPTI reconstruction under motion and *δ*B_0_ perturbations. Compared to the stationary reference, uncorrected data exhibit severe ringing artifacts, with NRMSE of 22.5% at TE = 10 ms and 35.4% at TE = 40 ms. vNAV-based correction (5-second temporal resolution) fails to improve image quality (NRMSE: 23.8% and 37.8%), again due to insufficient temporal resolution. In contrast, SMENA-based motion and *δ*B_0_ correction substantially improve image quality, reducing NRMSE to 11.0% and 15.7% at TE = 10 ms and 40 ms, respectively. Note that the stationary EPTI and the motion-corrupted EPTI are two separate scans with possible changes in position and background phases, which contribute to the residual errors after motion and *δ*B_0_ correction.

**Figure 9:**
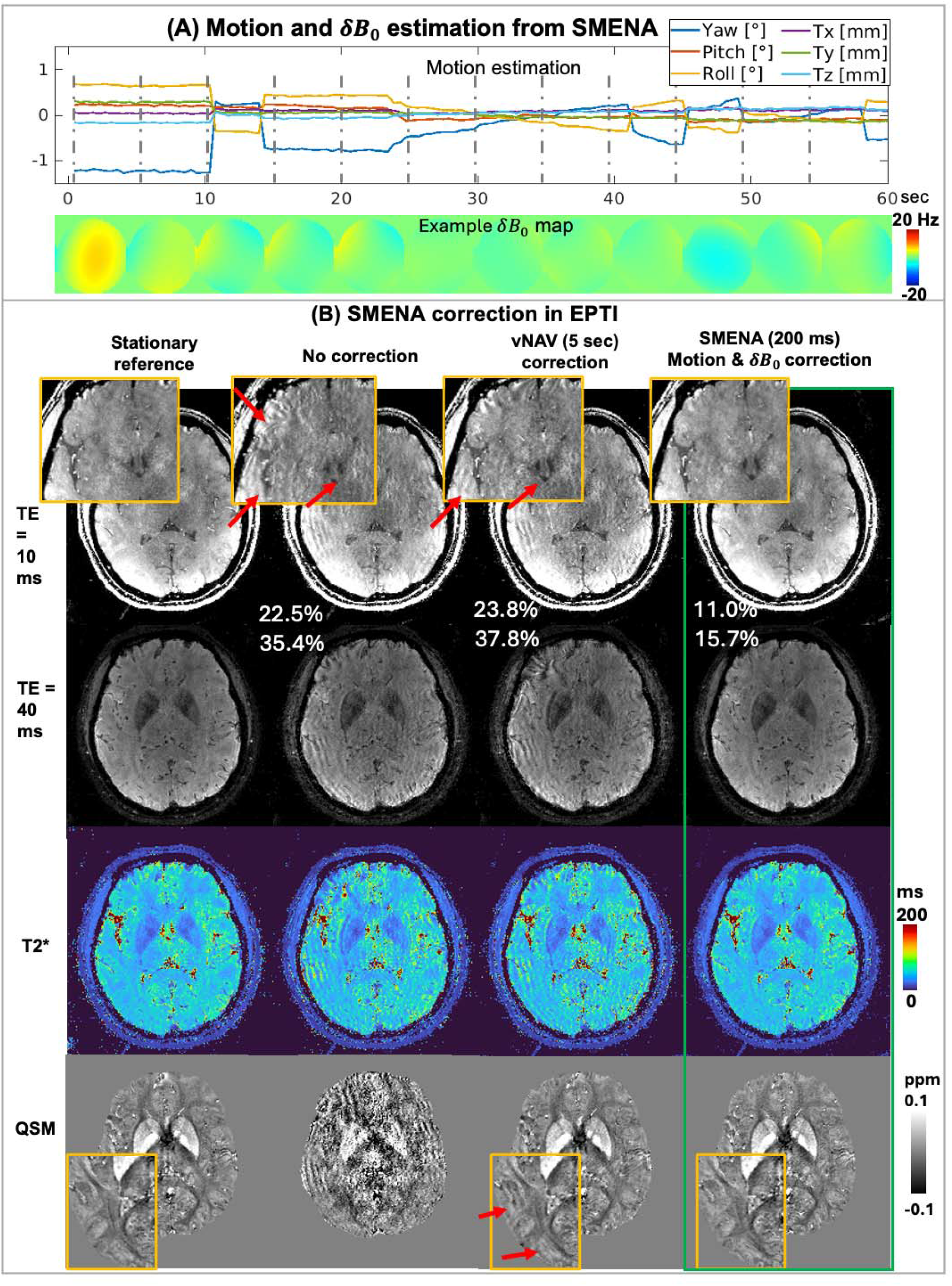
In vivo example of SMENA’s application to motion-corrupted GRE-EPTI. Similar to Figure 8, the correction with motion and *δ*B0 parameters from vNAV cannot provide good results due to the wrong timing, while the correction with SMENA’s motion and *δ*B0 parameters every 200 ms shows highly improved results, labeled by green frames.

Figure 10 presents an EPTI experiment with deep breathing. Although no intentional motion was introduced, the respiratory-induced motion, particularly pitch rotation, is evident (red line in Figure 10A). The average *δ*B_0_ across voxels closely follows the breathing pattern. Representative *δ*B_0_ maps, displayed at a 600-ms interval within a 7-second breathing cycle, further illustrate these dynamics. Without correction, noticeable shading artifacts are present in multi-echo images and derived maps. Motion-only correction using SMENA does not significantly improve image quality, indicating that *δ*B_0_ is the dominant source of artifacts in this scenario. Joint motion and *δ*B_0_ correction, in contrast, substantially improves quality of both images and quantitative maps.

**Figure 10:**
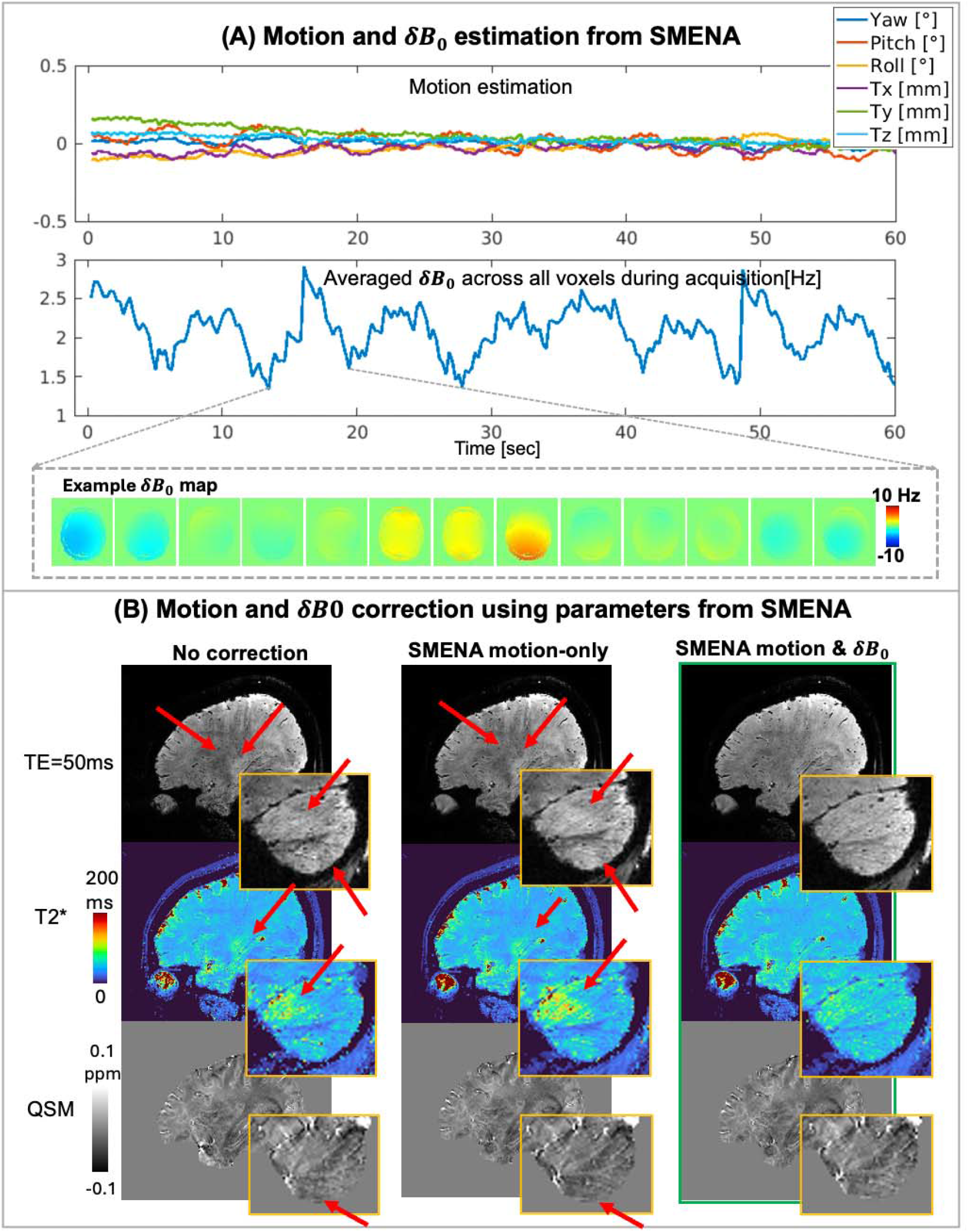
The application of SMENA on GRE-EPTI with deep breathing at 3T. In this case, the breathing-induced *δ*B0 changes drastic than the. SMENA, including motion parameters and *δ*B0, provides the best results (labeled by green frame). The highlighted artifacts are shown by red arrows.

Across simulations and in vivo experiments, SMENA consistently demonstrates that joint high-temporal-resolution estimation of motion and *δ*B_0_ is critical for accurate estimation and robust reconstruction. Additional examples with varying motion amplitudes are provided in Supporting Information Figure S4.

## 4. Discussions

Rigid motion and B0 perturbations have been longstanding challenges in MRI. Historically, motion artifacts and 0^th^-order *δ*B_0_ effect, which is also referred to as the global *δ*B_0_ effect, have been individually studied. In recent years, the high-order effect in *δ*B_0_ has been increasingly recognized, and the importance of jointly evaluating and correcting both motion and particularly *δ*B_0_ has been explored. Motion and *δ*B_0_ are often intrinsically coupled: rigid head motion—rotations in roll and pitch—can induce substantial *δ*B_0_ changes^41^, while physiological processes, such as deep breathing, can introduce strong *δ*B_0_ fluctuations and concomitant head motion. On the other hand, motion and *δ*B_0_ affect images through distinct mechanisms, and correcting only one of them is insufficient to fully restore image quality. Joint correction is therefore critical, particularly for long-TE and long-readout acquisitions such as multi-echo GRE and phase-sensitive contrasts such as SWI and QSM.

Accurate estimation of both motion and *δ*B_0_ is the foundation for addressing these challenges. Despite extensive prior work, it remains difficult to simultaneously satisfy several key requirements: (1) estimation with sub-second temporal resolution to capture both random and physiological variations; (2) joint estimation of motion and *δ*B_0_; (3) minimal additional scan time; and (4) ease of integration into existing clinical workflows. Recent developments such as Servo Navigator^27,28,42^ have demonstrated continuous estimation of motion and up to first-order*δ*B_0_; under conditions with small-to-moderate motion, but demonstration under large motion and higher-order*δ*B_0_; variations still needs further investigation.

In this work, SMENA addresses these challenges through a coordinated design of scout and navigator acquisitions that maximizes sensitivity to both motion and *δ*B_0_ while maintaining efficiency. The SMENA-scout employs an ultrafast multi-echo GRE-EPTI acquisition to obtain 3D 4-mm multi-echo images within 8 seconds. By leveraging temporal correlations along the echo dimension, the EPTI framework enables high acceleration while preserving image quality. Importantly, SMENA-scout provides T2* and *δ*B_0_ maps, allowing the generation of reference images at arbitrary echo times with matched magnitude and phase.

The compact navigator, SMENA-nav, also utilizes a multi-echo strategy with a carefully designed k-space trajectory to balance sensitivity to motion and *δ*B_0_. Specifically, the ky–kzsampling locations across echoes transition from central k-space (high sensitivity to *δ*B_0_), to higher-frequency k-space (high sensitivity to motion), and back to central k-space. Sampling central k-space at both early and late echoes enables accurate estimation of *δ*B_0_ through phase differences across echoes, which is more robust than single-echo phase-based estimation^13^. Paired with the contrast-matched reference images from SMENA-scout, motion and *δ*B_0_ become the only unknowns linking the predicted and acquired navigator signals, enabling robust joint estimation via the optimization framework described in Equation 2.

Both SMENA-scout and SMENA-nav employ similar 4-mm multi-echo Cartesian trajectories and gradient waveforms (Figure S1), leading to comparable short-term eddy current behavior. To further mitigate eddy current effects, field-correcting GRAPPA (FCG) ^38,39^ was applied to both SMENA-scout and SMENA-nav prior to motion and *δ*B_0_ estimation. In this work, SMENA-nav is placed at the beginning of each TR for simplicity. Alternative placements (e.g., during or after the main readout) may introduce additional eddy current effects accumulated from the main readout gradients. This can be addressed through FCG as well, or by performing one-time system calibration (e.g., using Skope^43^ or GIRF^44^).

Joint estimation of motion and *δ*B0 is critical for the accuracy of SMENA as well as other compact-navigator-based approaches. As demonstrated in Figure 6, neglecting *δ*B_0_ leads to Joint estimation of motion and *δ*B_0_ is critical for the accuracy of SMENA as well as other substantial errors in motion estimation, particularly for translations. This arises because translation induces linear phase modulation in k-space, while *δ*B_0_ introduces extra phase accumulation that manifests as a convolution kernel whose width increases approximately linearly with time. Under certain conditions, the effect of translation and *δ*B_0_ can mimic each other, leading to ambiguity in estimation. Joint modeling of motion and *δ*B_0_ effectively resolves this ambiguity and significantly improves accuracy.

Joint correction is also critical for restoring image quality. In both simulations (Figure 6) and in vivo experiments (Figure 8), motion-only correction leaves substantial residual artifacts, particularly in long-TE images where *δ*B_0_ effects are amplified. Even for short-TE images, joint Joint correction is also critical for restoring image quality. In both simulations (Figure 6) and in vivo experiments (Figure 8), motion-only correction leaves substantial residual artifacts, provides measurable benefits even in cases where *δ*B_0_ ‘s effects are less pronounced. correction consistently outperforms motion-only correction, indicating that *δ*B_0_ compensation provides measurable benefits even in cases where *δ*B0’s effects are less pronounced.

High temporal resolution estimation of motion and *δ*B_0_ is another key factor for effective correction. In the EPTI example (Figure 9), vNAV-based estimation at 5-second intervals fails to improve image quality due to incorrect timing of motion and *δ*B_0_ variations. Moreover, vNAV cannot adequately capture physiological *δ*B_0_ fluctuations, which typically oscillate periodically with a cycle of 1–4 seconds. In contrast, SMENA achieves a temporal resolution of ∼200 ms, enabling accurate tracking of both random motion and respiration-induced variations. This capability is particularly valuable for motion-prone populations such as pediatric and geriatric subjects.

Incorporating high-temporal-resolution motion and *δ*B_0_ parameters from SMENA into the image reconstruction introduces increased computational demands, as the number of Fourier transform and inverse Fourier transform grows linearly with the number of motion/ *δ*B_0_ states, which can end up in thousands. While clustering the motion and *δ*B_0_ states into fewer groups can reduce computational burden, they may be less effective in scenarios with rapid or large-amplitude motion. To address this, our group has developed a Multi-Layer Perceptron (MLP)-based reconstruction method, mobile-GRAPPA^45^, which reduces the problem to a single-state reconstruction while preserving high-temporal-resolution motion/ *δ*B_0_ correction. This approach provides a promising direction for scaling SMENA to more challenging applications.

Despite the substantial improvements achieved by SMENA on motion/ *δ*B_0_ corrupted data, the corrected image quality remains slightly inferior to that of fully stationary acquisitions, even in simulations. One contributing factor is that motion alters the effective k-space sampling distribution, which degrades the conditioning of the reconstruction problem. This observation motivates the integration of prospective motion correction to further improve image fidelity. Additionally, the current implementation relies on a quasi-Newton optimization algorithm, requiring approximately 1 minute per state for parameter estimation. Future work will focus on accelerating this process. Recent studies have demonstrated near real-time estimation of motion and low-order *δ*B_0_ using linear regression approaches^27,28^, which could be integrated into the SMENA framework.

One limitation of SMENA is the acquisition time of the SMENA-scout. Although 5-8 seconds is already short, it can be problematic for motion-prone populations such as infants. A potential solution is to re-order the sampling pattern for SMENA-scout to achieve self-navigation^24^. Another limitation is that *δ*B_0_ is modeled using a global second-order spherical harmonic representation due to the limited navigator data. At 3T and under moderate motion or breathing, this approximation is generally sufficient. However, under ultra-large motion (e.g., >10° rotation) or at ultra-high field strengths (≥7T), higher-order spatial variations in *δ*B_0_ become significant and cannot be captured by a second-order model. Future work will focus on exploiting spatiotemporal redundancy in *δ*B_0_ to enable higher-fidelity estimation with increased spatial complexity.

## 5. Conclusion

SMENA enables accurate, high-temporal-resolution estimation of motion and *δ*B_0_ with minimal scan-time overhead. When integrated into multi-echo GRE and GRE-EPTI acquisitions, SMENA substantially improves image quality and quantitative mapping under conditions of random motion and deep breathing. Joint estimation of motion and *δ*B_0_ is critical for estimation accuracy, while joint correction is essential for high-quality image reconstruction. Future studies in motion-prone populations are warranted to further evaluate its clinical utility.

## 6. Data availability statement

The example data for SMENA pipeline are publicly available at: https://github.com/nanwangpku/Motion_estimation.git

## Figure Caption

**Supporting Information Figure S1:** The gradient waveforms for SMENA-nav and SMENA-scout. Both gradient waveforms use the same maximum gradient and slew rate, inducing similar structure. Maximum ramp sampling is used to allow the most efficient sampling under hardware limits.

**Supporting Information Figure S2:** The Order of the motion operator and phase operator matters. (A) when directly performing motion and phase operators to a static image, which is equal to moving an image to a different head position, performing the motion operator separately on the magnitude and phase of the image produced more dotting artifacts than performing the motion operator once after combing the magnitude and phase of the image. (2) In an actual reconstruction of GRE-EPTI data, this effect is elevated. When performing motion operator separately to the image to the resolved and the phase, it produced large dotting and streaking errors compared to performing the motion operator after the combination of image and its phase.

**Supporting Information Figure S3:** The comparison of motion-only estimation and joint estimation of a in vivo case. When doing motion-only estimation, SMENA’s results and vNAV’s results at the 12 time points shows a large discrepancy, especially for translation. The motion parameters using joint motion and dB0 estimation shows a reduced error compared to vNAV.

**Supporting Information Figure S4:** Two more in vivo examples

