## Supporting Information for "Scout-based Multi-Echo NAvigator (SMENA) for high temporal resolution motion and B_0_ estimation and correction: applications to multi-echo GRE and EPTI"

A. **Gradient waveform design of SMENA**

The gradient waveforms of SMENA-nav are shown in Figure S1 A. The maximum slew rate is 120 mT/m/ms; the maximum gradient for 4-mm resolution is 26 mT/m. The maximum ramp sampling portion is used to minimize the echo spacing as well as the add-on time for SMENA-nav.

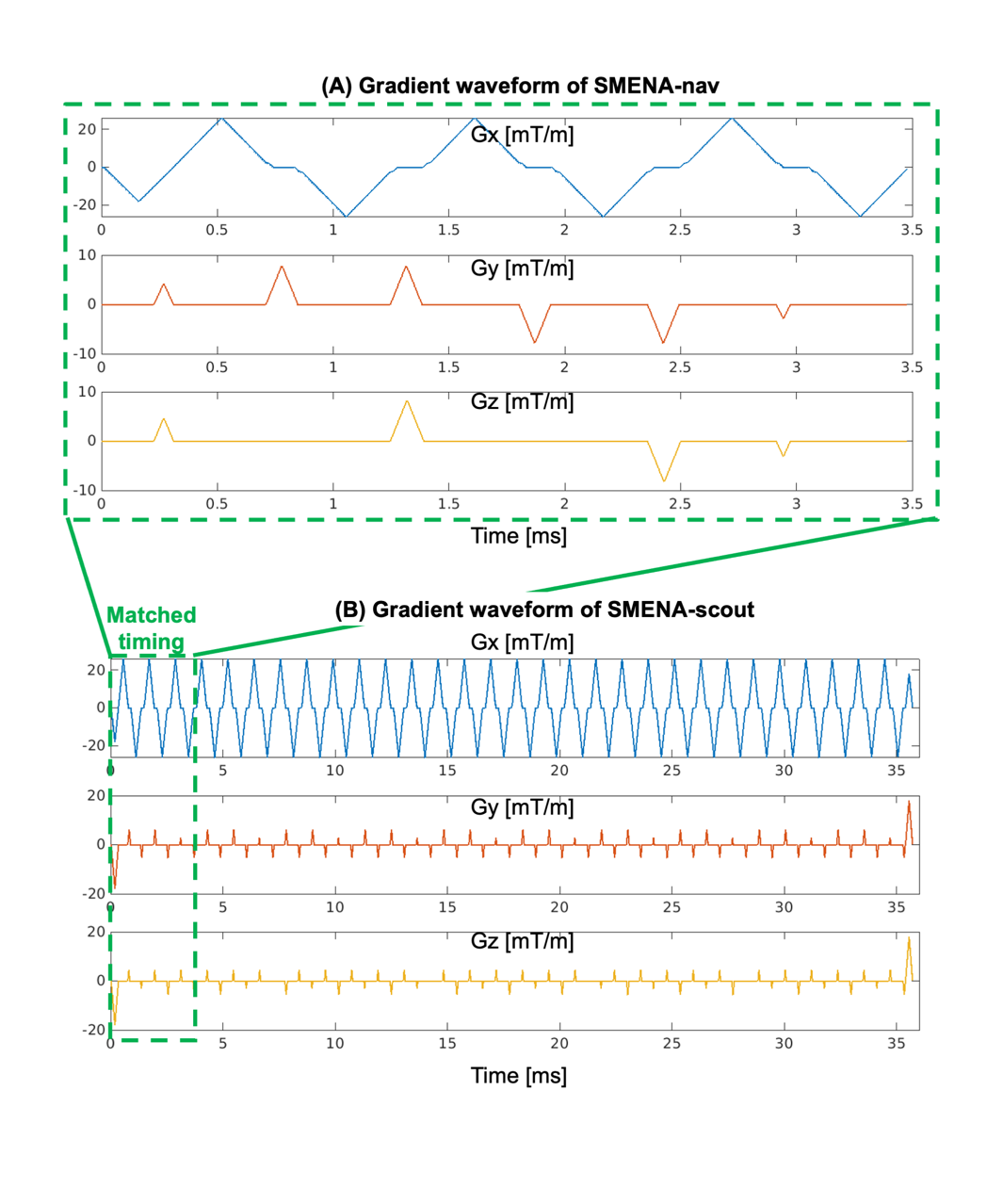

**Figure S1:** The gradient waveforms for SMENA-nav and SMENA-scout. Both gradient waveforms use the same maximum gradient and slew rate, inducing similar structure. Maximum ramp smapling is used to allow the most efficient sampling under hardware limits.

Because the SMENA-nav is acquired with full ramp sampling, the spatial-temporal varying phase errors related to eddy current from changing gradient can be an issue. In this work, the issue is alleviated by two approaches. First, a similar gradient waveform pattern is used for SMENA-scout, with one example shot shown in Figure S1 B. The maximum slew rate, maximum gradient, and portion of ramp sampling in SMENA-scout matched with SMENA-nav. The similarity in the gradient waveform designs induces similarity in the eddy current patterns.

Second, FCG technique, which was developed by our group to minimize the eddy-current-induced errors for EPI-based readout, has been implemented as a pre-processing step to correct the SMENA-scout and SMENA-nav data. To perform FCG, two shots of SMENA-scout and SMENA-nav are acquired with opposite readout polarity. A set of MLP-based kernels is trained to correct the single-polarity data to the dual-polarity averaged data, which is considered as eddy-current free. The trained kernels are then used to correct the actual SMENA-scout and SMENA-nav data before the image reconstruction. These two approaches substantially reduce eddy-current-induced phase errors and improve the consistency between the scout images and navigators, ensuring accurate motion and $\delta B_{0}$ estimation.

B. **Permutation of motion and phase operator**

Determining the correct order of the operator for B0-induced phase and the operator for motion is important for the estimation and correction of motion and $\delta B_{0}$. Mathematically, there are two ways to define the operators of motion and B0-induced phase:

$$I_{\boldsymbol{\theta,}\delta B_{0}}=\mathbf{T}\mathbf{P}\left( B_{0}\boldsymbol{+}\delta B_{0} \right)\boldsymbol{I}_{\boldsymbol{0}} (S.1)$$

or

$$I_{\boldsymbol{\theta,}\delta B_{0}}=\mathbf{T}\boldsymbol{(}\mathbf{P}\left( B_{0}\boldsymbol{+}\delta B_{0} \right)\boldsymbol{)}{\mathbf{T}\boldsymbol{(I}}_{\boldsymbol{0}}\boldsymbol{)}(S.2)$$

Equation S.1 models the production of phase and real-valued images at the original position first, followed by the motion operator on the complex-valued images. In contrast, Equation S.2 first rotates the phase map and the real-valued images to the target position separately, followed by the production of them to obtain the complex-valued images. The motion in the image space, when there is rotation, operates as circular convolution in the k-space domain. In practice, since the images are acquired with finite resolution, the circular convolution induces errors at the k-space edge. Therefore, given a finite-resolution real-valued image and a corresponding phase map, Equation S.1 and S.2 are not equal, as shown in Figure S2 A. The images created with Equation S.2 bear more high-resolution errors, while results with Equation S.1 show clean outcomes.

The performance of both models is further demonstrated in the reconstruction of motion-corrupted images, as shown in Figure S2 B. The image reconstructed using the imaging model described by Equation S.1 displays cleaner structure and less flickering. Therefore, the order of motion operator and phase operator, for both estimation (Equation 2) and reconstruction (Equation 5 & 6), is determined based on Equation S.1.

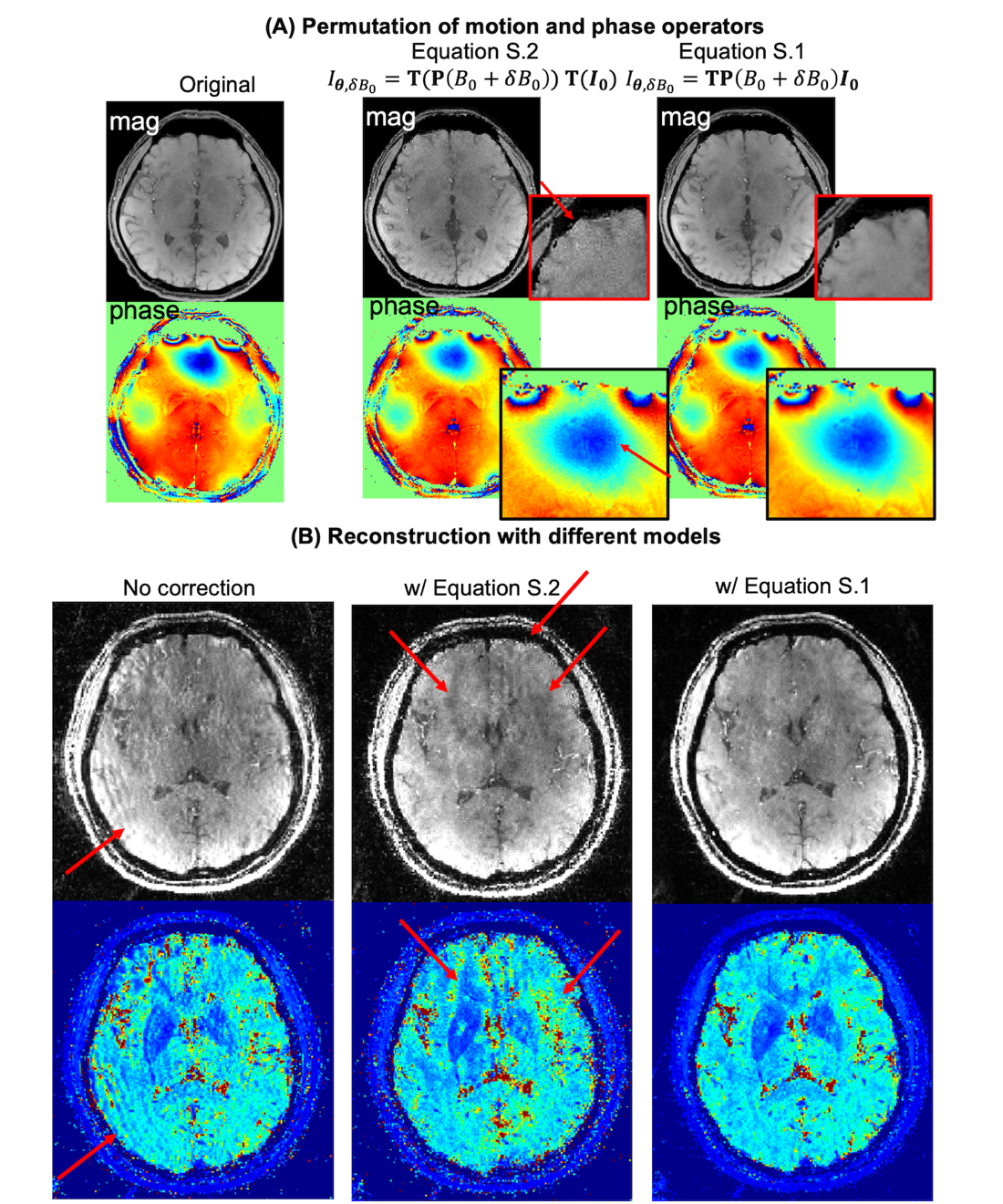

**Figure S2:** The Order of the motion operator and phase operator matters. (A) when directly performing motion and phase operators to a static image, which is equal to moving an image to a different head position, performing the motion operator separately on the magnitude and phase of the image produced more dotting artifacts than performing the motion operator once after combing the magnitude and phase of the image. (2) In an actual reconstruction of GRE-EPTI data, this effect is elevated. When performing motion operator separately to the image to the resolved and the phase, it produced large dotting and streaking errors compared to performing the motion operator after the combination of image and its phase.

C. **Superiority of joint motion and** $\boldsymbol{\delta}\boldsymbol{B}_{\boldsymbol{0}}$ **estimation**

The significance of performing joint motion and $\delta B_{0}$ estimation has been evaluated in the Simulations, with the results showing that joint estimation provides substantially improved accuracy. The in vivo results show consistent indications, as illustrated in Figure S3. The results are from the same in vivo study as shown in Figure 7. When performing motion-only estimation with SMENA-nav, the errors between SMENA estimation and vNAV estimation at the vNAV time points are elevated (Figure S3 A), particularly for the estimation of translation; with joint motion-and- $\delta B_{0}$-estimation, the error is largely reduced (Figure S3 B).

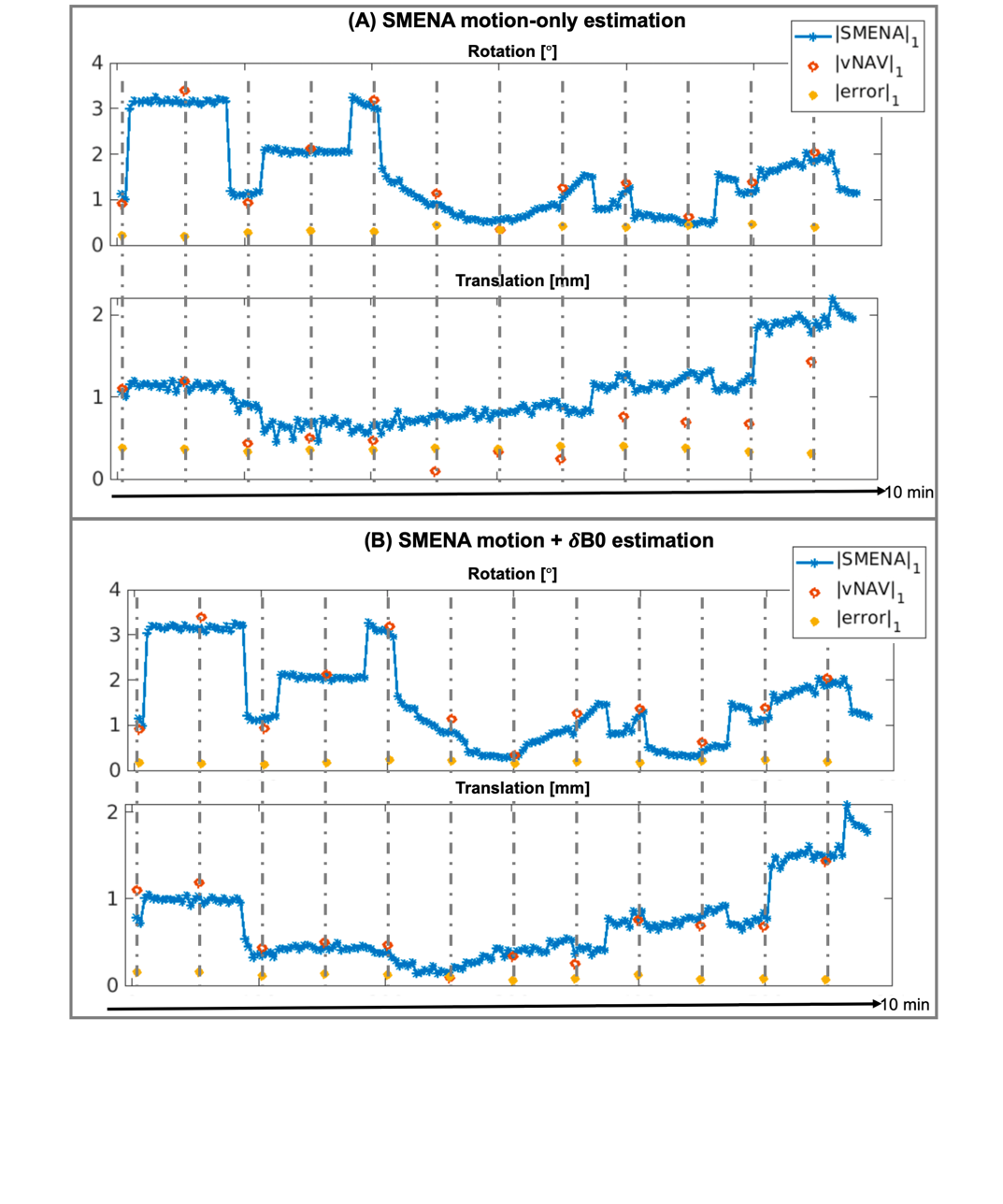

**Figure S3:** The comparison of motion-only estimation and joint estimation of a in vivo case. When doing motion-only estimation, SMENA’s results and vNAV’s results at the 12 time points shows a large discrepancy, especially for translation. The motion parameters using joint motion and dB0 estimation shows a reduced error compared to vNAV.

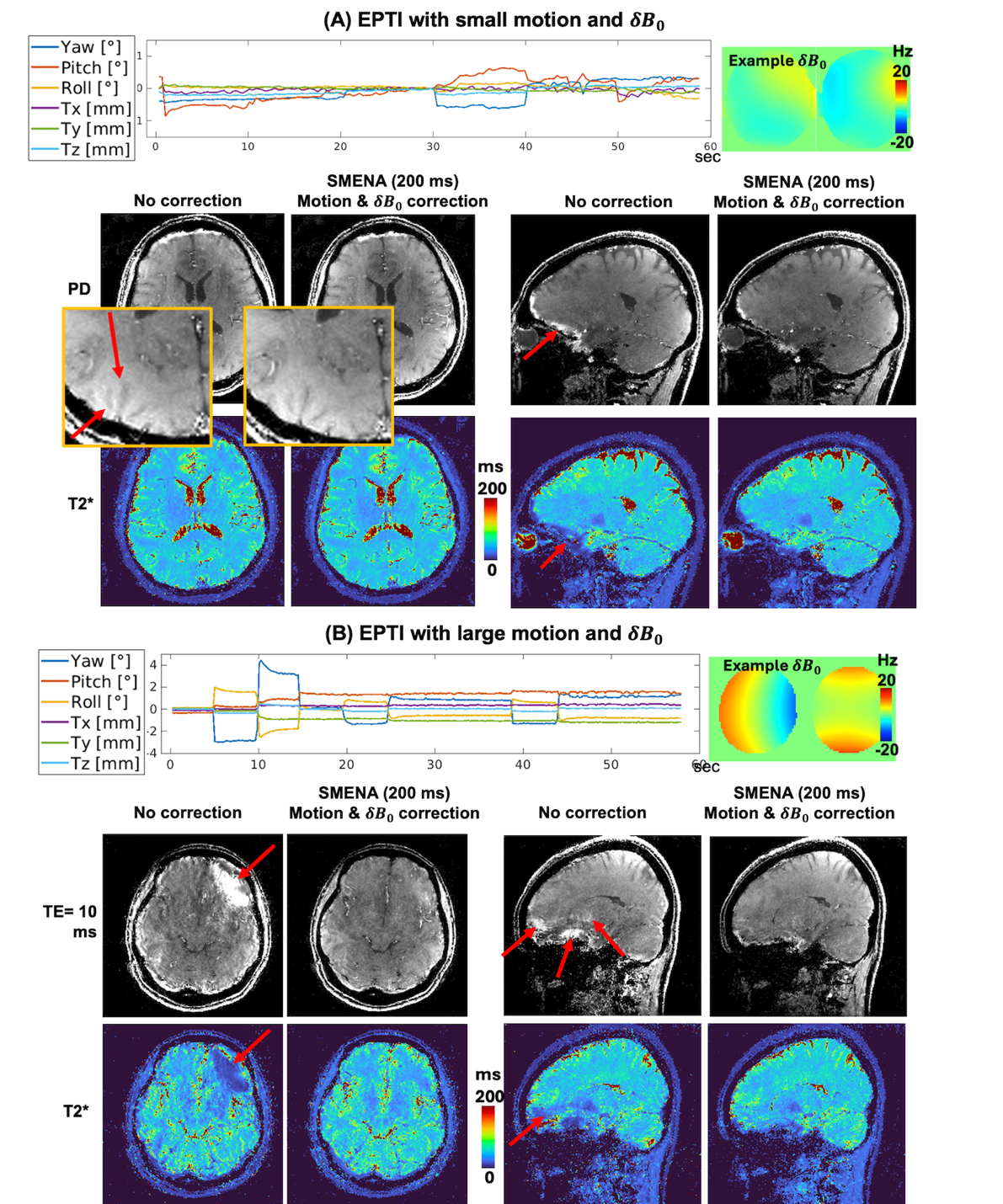

**Figure S4:** Two more in vivo examples.

D. **Additional in vivo examples with SMENA**

Two additional in vivo GRE-EPTI examples are demonstrated in Figure S4. An example with small motion is displayed in (A). The results with SMENA motion- and- $\delta B_{0}$-correction can further sharpen the structure and reduce the illumination from B_0_ errors. For another study with large and multi-state motion (Figure S4 B), the images without any correction bear significant artifacts, while the image quality with joint motion- and- $\delta B_{0}$ correction is largely restored.
